# Host genome integration and giant virus-induced reactivation of the virophage mavirus

**DOI:** 10.1101/068312

**Authors:** Matthias G. Fischer, Thomas Hackl

**Affiliations:** Department of Biomolecular Mechanisms, Max Planck Institute for Medical Research, 69120 Heidelberg, Germany

## Abstract

Endogenous viral elements (EVEs) are increasingly found in eukaryotic genomes^1^, yet little is known about their origins, dynamics, or function. Here, we provide a compelling example of a DNA virus that readily integrates into a eukaryotic genome where it acts as an inducible antiviral defense system. We found that the virophage mavirus^2^, a parasite of the giant Cafeteria roenbergensis virus (CroV)^3^, integrates at multiple sites within the nuclear genome of the marine protozoan *Cafeteria roenbergensis*. The endogenous mavirus is structurally and genetically similar to the eukaryotic Maverick/Polinton DNA transposons^4,5^ and endogenous polintoviruses^6^. Provirophage genes are not constitutively expressed, but are specifically activated by superinfection with CroV, which induces the production of infectious mavirus particles. Virophages inhibit the replication of giant viruses and a beneficial effect of provirophages on their host cells has been hypothesized^2,7^. We found that provirophage-carrying cells are not directly protected from CroV; however, lysis of these cells releases reactivated mavirus particles that are then able to suppress CroV replication and enhance host survival of other CroV-infected flagellate populations in a dose-dependent manner. The host-parasite interaction described here involves an altruistic aspect that is unique among microbes. Our results demonstrate a direct link between mavirus and Maverick/Polinton elements and suggest that provirophages can defend natural protist populations against infection by giant viruses.

---

Virophages of the family *Lavidaviridae*^8^ are obligate parasites of giant DNA viruses. During co-infection of a protist host, these 15-30 kbp dsDNA viruses replicate in the cytoplasmic virion factory of the giant virus, which results in decreased giant viral progeny and increased host survival rates^2,9^. Mavirus-like virophages share a common evolutionary origin with a class of self-synthesizing DNA transposons called Maverick/Polinton elements (MPEs)^4,5^. MPEs are widespread in eukaryotic genomes and encode virus-like genes, which led to their recent designation as "polintoviruses", representing perhaps the most broadly distributed family of EVEs among eukaryotes^6,10,11^. Virophages encode integrase genes, and endogenous virus genomes (so-called provirophages) have been reported in mimivirus^12^ and in the nuclear genome of the marine alga *Bigelowiella natans*^7^. Endogenous virophages in protists are hypothesized to protect the host cells from giant virus infection^2,7^, but experimental data for this claim were lacking.

To generate host-integrated provirophages, we infected the clonal *C. roenbergensis* strain E4-10P with the giant virus CroV at a multiplicity of infection (MOI) of 0.01 and with mavirus at MOI ≈1 and screened the surviving cells for mavirus DNA (Fig. 1a, Extended Data Fig. 1). PCR analysis of 66 clonal survivor strains identified 21 (32%) mavirus-positive host strains. We chose the cell line with the strongest mavirus signal for further analysis and named it E4-10M1. Filtration of E4-10P and E4-10M1 cell populations through syringe filters of various pore sizes and qPCR analysis of the resulting filtrates confirmed that the observed mavirus signal was associated with E4-10M1 cells and not caused by remaining free virus particles (Extended Data Table 1). We then sequenced genomic DNA from strains E4-10P and E4-10M1 on Illumina MiSeq and Pacific BioSciences (PacBio) RS II platforms and created hybrid assemblies for each strain. The sequence data suggested that *C. roenbergensis* has a diploid genome (Extended Data Figure 2), which obstructed the direct assembly of mavirus-containing contigs because integration at a specific site occurred at only one of the two alleles, thus introducing a structural ambiguity that led to erroneous assemblies. We therefore scanned the E4-10M1 genome assembly indirectly for integrated mavirus sequences by aligning corrected PacBio reads to the mavirus reference genome, extracting those reads, and assembling them into contigs. The longest resulting contig was 30,556 bp in length and contained a 19,055 bp sequence that was 100% identical to the mavirus reference genome (GenBank accession HQ712116). In contrast to the reference mavirus genome, the endogenous virus genome was flanked on either side by 615/616 bp-long TIRs that were 99.7% identical to each other. The longer TIRs result in a total length of 20,190 bp for the endogenous mavirus genome, compared to 19,063 bp for the reference genome. By recruiting reads to the flanking regions of mavirus TIRs, we found 11 well-supported integration sites in the E4-10M1 genome (Extended Data Table 2). The host sequence directly adjacent to the provirophage genome featured target site duplications (TSDs) that were 5-6 bp long. The TSD sequences differed between integration sites with no obvious consensus motif. One of the integration sites was characterized in further detail and its reconstruction is shown in Fig. 1b. PCR analysis verified the predicted integration site and confirmed that the E4-10M1 strain is heterozygous for the integrated mavirus genome (Fig. 1c+d).

**Figure 1:**
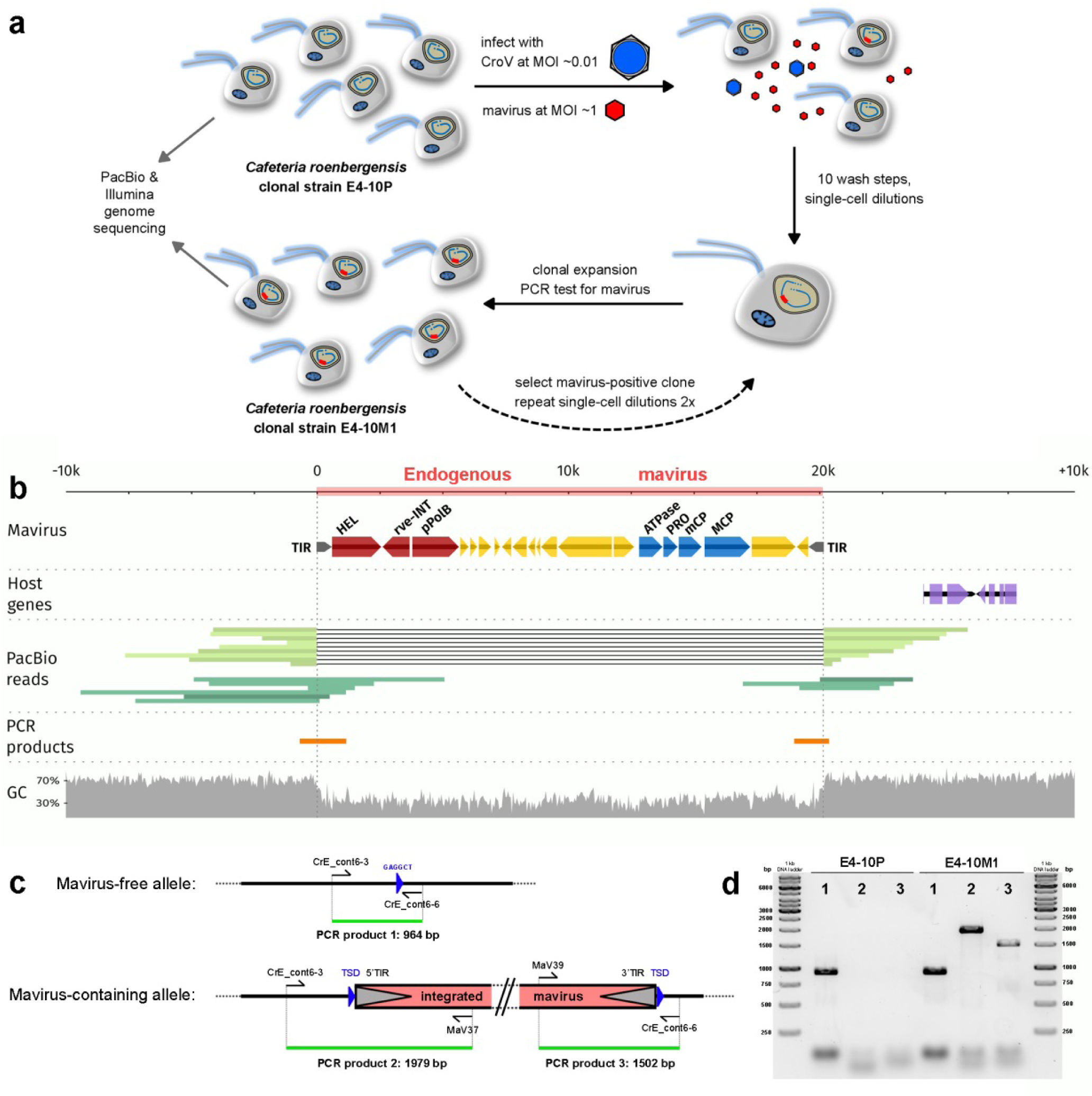
Generation and characterization of endogenous mavirus virophages in Cafeteria roenbergensis. **a**, The clonal C. roenbergensis *strain E4-10P was infected with CroV and mavirus and surviving cells were PCR-tested for mavirus. The genomes of the resulting mavirus-positive clonal strain E4-10M1 and of the parental E4-10P strain were sequenced on PacBio and Illumina platforms to analyze mavirus integration sites. **b**, Partial view of a 208 kbp E4-10M1 contig featuring the 20,190 bp long mavirus genome flanked by 10 kbp of host sequence on either side. Mavirus genes for replication and integration are shown in red, morphogenetic genes are shown in blue, other genes are shown in yellow and terminal inverted repeats (TIR) are indicated in grey. The exon structure of two adjacent host gene models (function unknown) is shown in purple. PacBio reads covering the integration site are shown in green. Reads that span the integration site and contain only host sequence are shown in light green, whereas reads that cross the virus-host junction are shown in dark green. PCR products spanning the host-virus junction are shown in orange. The GC content plot is based on a 30 bp sliding window. **c**, Schematic representation of the host genomic region in b illustrating the PCR primer binding sites and expected PCR products that were used to confirm the integration site. **d**, Gel image of the PCR products obtained from E4-10P and E4-10M1 genomic templates with primers spanning the integration site.The lanes are labelled according to the primer combinations and products shown in c. L, DNA ladder.*

To test if the endogenous mavirus genes were expressed, we analyzed selected transcripts by reverse transcription qPCR. Mavirus gene promoters are highly similar to the late gene promoter motif in CroV^2^, which suggests that CroV can activate mavirus genes. We therefore isolated total RNA from mock-infected and CroV-infected E4-10P and E4-10M1 cells at 0 and 24 h p.i. and quantified in the resulting cDNA pool five mavirus genes, three CroV genes, and one host cell gene using gene-specific primers (Extended Data Table 3). CroV transcripts could be clearly detected at 24 h p.i. in the infected cultures and their expression levels were comparable between E4-10P and E4-10M1 strains. The mavirus genes in E4-10M1 cells were quiescent under normal conditions and also immediately after inoculation with CroV. However, mavirus genes were strongly expressed at 24 h p.i. in the CroV-infected E4-10M1 strain (Fig. 2). CroV infection thus induces expression of the endogenous mavirus genes in E4-10M1 cells. Addition of the protein biosynthesis inhibitor cycloheximide (CHX) or the DNA polymerase inhibitor aphidicolin (APH) effectively inhibited host cell growth and CroV DNA replication (Supplemental Spreadsheet). CHX treatment inhibited expression of the intermediate DNA polymerase B gene *crov497* and of the late major capsid gene *crov342*, but not of the early isoleucyl-tRNA synthetase gene *crov505* (Extended Data Fig. 3). This is in line with the presence of a viral transcription apparatus in the virion of CroV^13^ and other cytoplasmic large DNA viruses^14,15^, which mediates early viral gene expression. In the presence of APH, all three CroV genes were expressed at low levels, with cDNA of the late MCP gene being barely detectable. Crucially, treatment with CHX or APH also inhibited mavirus gene expression in CroV-infected E4-10M1 cells, indicating that *de novo* protein synthesis and CroV DNA replication are prerequisites for provirophage gene induction. Based on these results and the similarity of transcriptional signals between virophages and their giant host viruses, we propose that a CroV-encoded late transcription factor may be responsible for provirophage activation (Extended Data Fig. 4).

**Figure 2:**
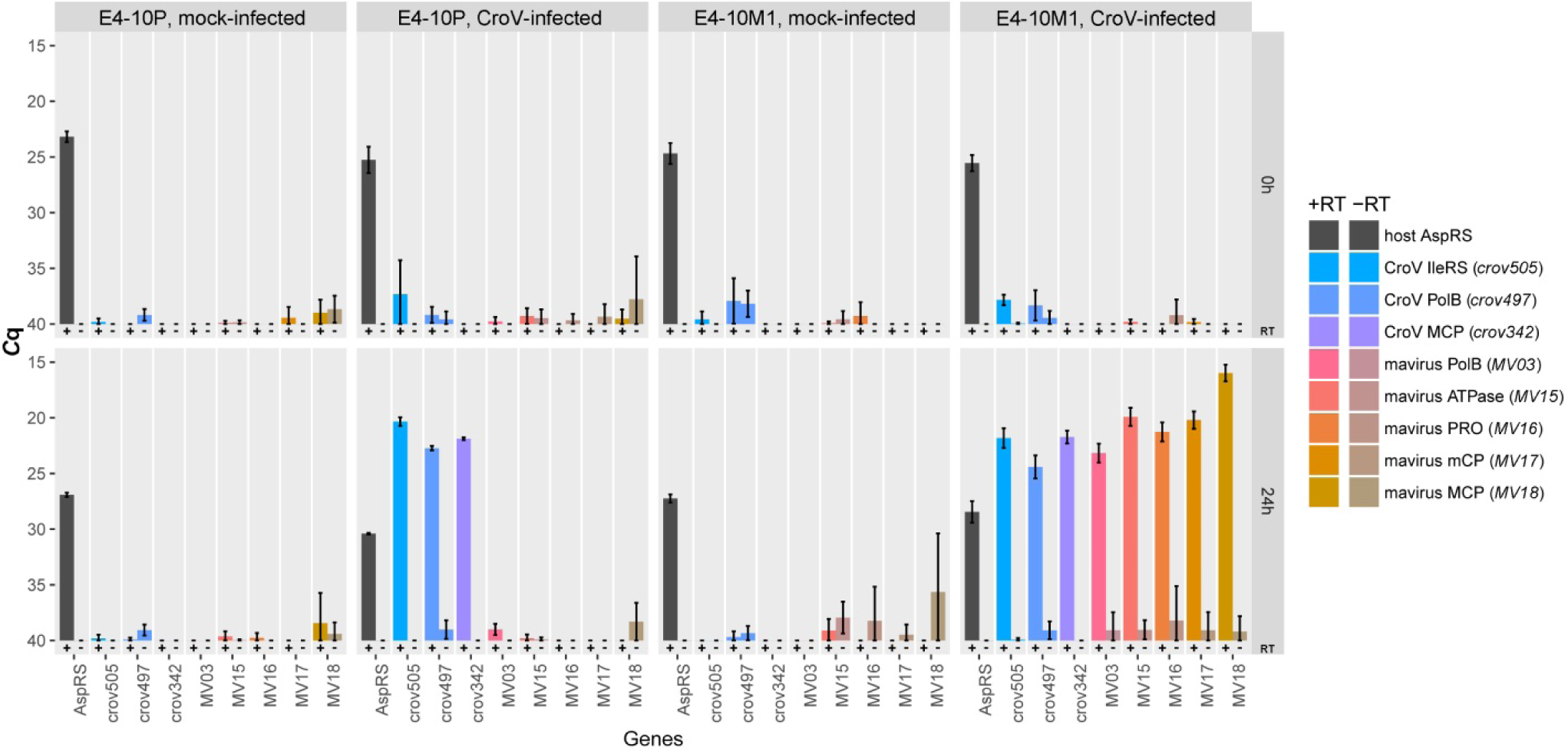
Gene expression analysis of the endogenous mavirus genome. *Selected cellular and viral transcripts isolated at 0 and 24 h p.i. from mock-infected or CroV-infected E4-10P and E4-10M1 cultures were quantified by qRT-PCR. Shown are the average quantification cycle (Cq) values of three independent experiments with error bars representing ± SD. The following genes were assayed: host AspRS,* C. roenbergensis *E4-10 aspartyl-tRNA synthetase;* crov342*, CroV major capsid protein;* crov497*, CroV DNA polymerase B;* crov505*, CroV isoleucyl-tRNA synthetase;* MV03*, mavirus DNA polymerase B;* MV15*, mavirus genome-packaging ATPase;* MV16*, mavirus maturation protease;* MV17*, mavirus minor capsid protein;* MV18*, mavirus major capsid protein. Cq values of the reverse transcriptase (RT) negative reactions are shown directly to the right of the respective RT positive results. Accession numbers are listed in Extended Data Table 3. See also Supplemental Spreadsheet.*

Next, we examined whether CroV infection would induce DNA replication of the integrated mavirus genome. E4-10P and E4-10M1 cells were either mock-infected or CroV-infected and viral DNA levels were monitored by qPCR. No virus DNA was found in mock-infected E4-10P cells, whereas a latent mavirus signal was present in mock-infected E4-10M1 cells (Fig. 3a). In CroV-infected E4-10M1 cells, the mavirus signal increased ≈500fold within 48 h p.i., proving that CroV induces genome replication of the mavirus provirophages. CroV replication and cell lysis were comparable in both host strains and also the CroV titer was similar in E4-10P and E4-10M1 lysates (≈5E+07 per mL), indicating that provirophage induction does not inhibit CroV propagation or prevent cell lysis in CroV-infected E4-10M1 cells. Electron microscopy of concentrated and purified cell lysates revealed the presence of mavirus-like particles in CroV-infected E4-10M1, but not E4-10P lysates, nor in mock-infected cultures (Fig. 3b+c, Extended Data Fig. 5). To test whether these particles are infectious, we co-inoculated E4-10P cultures with 0.1 µm-filtered material from mock- or CroV-infected E4-10P and E4-10M1 cultures (Fig. 3d). Only lysate from CroV-infected E4-10M1 cultures contained mavirus DNA, which replicated in the presence of CroV. We conclude from these results that CroV induces the production of infectious mavirus particles in strain E4-10M1. Interestingly, the reactivated mavirus suppressed CroV genome replication by 2-3 orders of magnitude, resulting in survival of the host cell population (Fig. 3d). Treatment of reactivated mavirus with 500 J/m^2^ of ultraviolet light (λ=254 nm) prior to infection abrogated these effects (Extended Data Fig. 6), suggesting that mavirus is indeed the causative agent.

**Figure 3:**
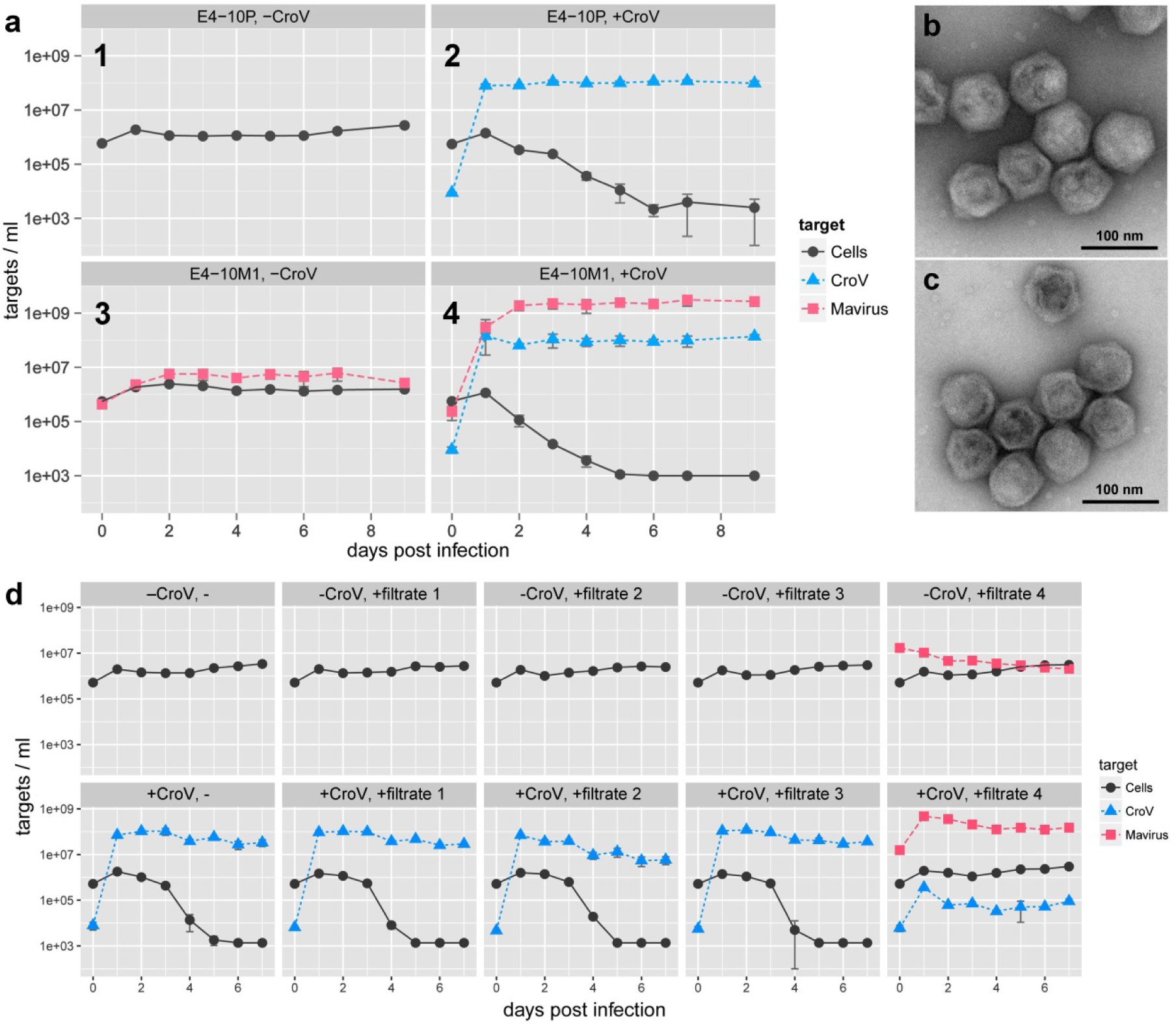
CroV infection induces replication and virion production of the endogenous mavirus. ***a**, C. roenbergensis strains E4-10P and E4-10M1 were mock-infected (1,3) or infected with CroV (2,4) and cells and viruses were monitored for 9 days. Cell densities are based on microscopy counts, virus concentrations are derived from qPCR data. **b**, Negative-stain electron micrograph of virus particles from the CroV-infected E4-10M1 strain (panel 4 in **a**). **c**, Electron micrograph of reference mavirus particles. **d**, Strain E4-10P was mock-infected (upper row) or CroV-infected (lower row) and simultaneously inoculated with 0.02% (v/v) of 0.1 µm filtrates sampled at 3 d p.i. during the infection experiments 1-4 shown in **a**). Data in **a**) and **d**) were pooled from three independent experiments and error bars represent ± SD. See also Supplemental Spreadsheet.*

To gain more insight in the virophage-virus-host dynamics, we infected E4-10P cells with different MOIs of CroV and of reactivated mavirus. Figure 4a shows infections with CroV MOIs of 0.01 to 10 in the absence or presence of mavirus at MOI ≈10. The number of virions that each cell receives at a given MOI follows a Poisson distribution, therefore the percentage of infected cells at an MOI of 1 is 63%, and an MOI of 10 is needed to ensure that >99.99% of cells are infected. With every cell infected with mavirus, host populations survived an infection with CroV at MOIs of 0.01 to 1 (Fig. 4a). Although CroV at MOI 10 did not replicate in the presence of mavirus, the cells still lysed (96.5% decline after 5 days). These data indicate that nearly every cell infected with CroV dies, irrespective of mavirus, and that mavirus rather halts the spread of CroV by inhibiting its replication and preventing the release of progeny virions from lysed co-infected cells. When E4-10P cells were infected with a CroV MOI of 1 and mavirus MOIs ranging from ≈0.001 to ≈10, a clear dose-response relationship was observed for host survival and inhibition of CroV DNA replication (Fig. 4b). Even low MOIs of mavirus significantly inhibited CroV and improved host survival rates (Fig. 4c).

**Figure 4:**
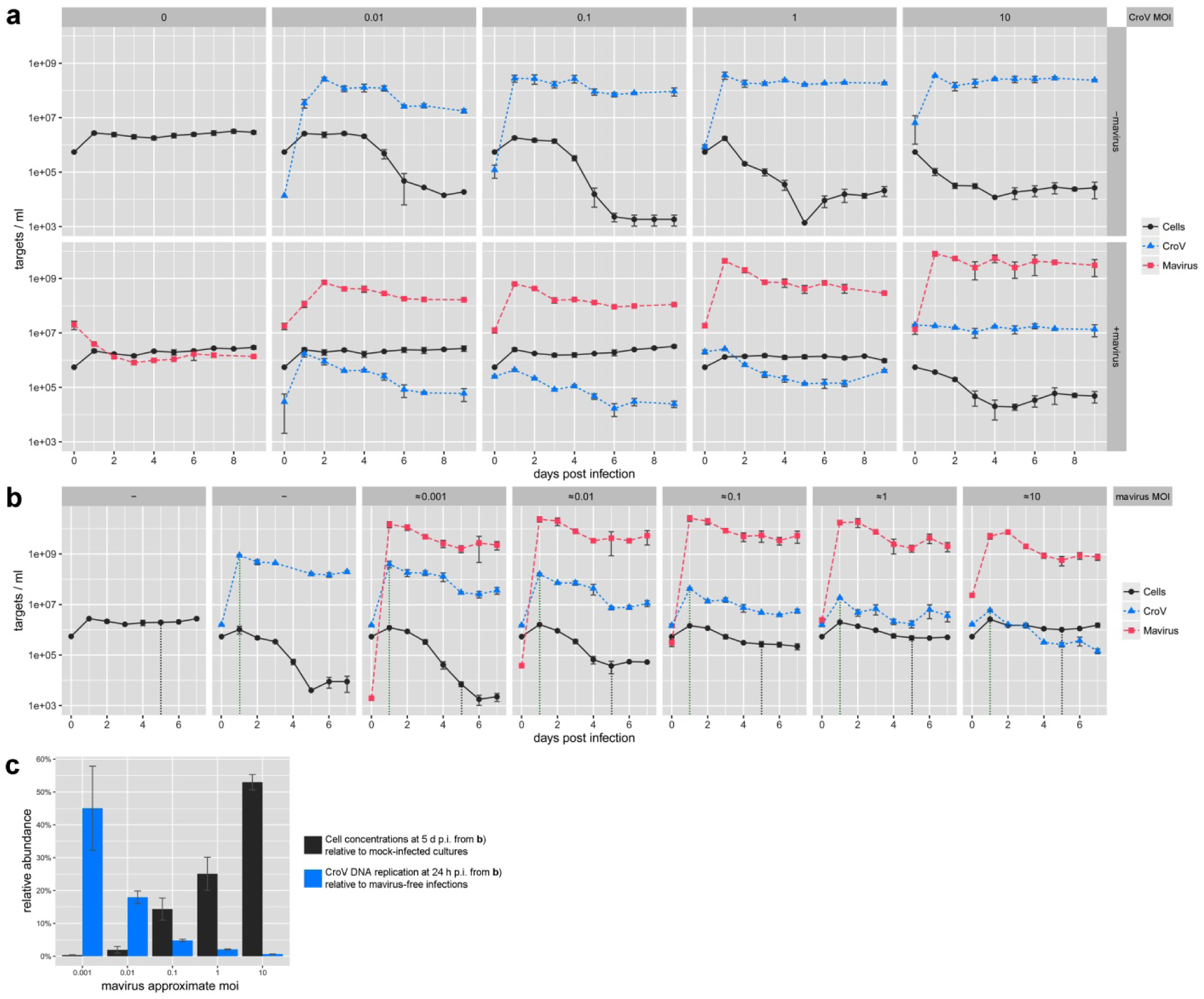
Reactivated mavirus inhibits CroV and promotes host survival in subsequent infections. C. roenbergensis *strain E4-10P were infected with different MOIs of CroV and mavirus. Mavirus inocula represent the 0.1 µm filtrate of the CroV-infected E4-10M1 culture at 3 d p.i. (panel 4 in Fig. 3a). Cell densities are based on microscopy counts, virus concentrations are derived from qPCR data. Shown are the average values of biological triplicates and error bars represent ± SD. See also Supplemental Spreadsheet. **a**, Infection experiments with different MOIs of CroV in the absence (upper row) or presence (lower row) of mavirus at an MOI of ≈10. **b**, Cultures of* C. roenbergensis *strain E4-10P infected with CroV at MOI=1 and increasing MOIs of reactivated mavirus. The leftmost panel shows the mock-infected control, the second panel from the left shows the mavirus-free, CroV-infected control. Vertical dotted lines mark the reference points for the analysis in **c**). **c**, Summary of the effect of mavirus MOI on CroV DNA replication and host cell survival from the infection experiments shown in b. Black columns show the host cell densities at 5 d p.i. with increasing mavirus MOIs relative to mock-infected 5 d p.i. cultures. Blue columns show the CroV genome copy concentration at 24 h p.i. with increasing mavirus MOIs relative to mavirus-free 24 h p.i. CroV infections.*

Our results show that mavirus readily integrates in the nuclear genome of its eukaryotic host *C. roenbergensis* and that mavirus provirophages resemble MPEs not only in length, gene content and host range, but also in their TIR and TSD structures. We demonstrate that endogenous mavirus genes are transcriptionally silent unless the cell is infected with CroV, which triggers gene expression, genome replication, and virion synthesis of the provirophage. Although provirophage-carrying cells are not directly protected from CroV, lysis of these cells releases infectious mavirus particles that are able to inhibit CroV in subsequent co-infections. Provirophagemediated host defense against giant viruses in the *Cafeteria* system thus follows an altruistic model, in which some cells are sacrificed in order to protect their kin. Our study also reveals a symbiotic virus-host relationship, in which the cell provides an opportunity for mavirus to persist as a provirophage while the host population benefits from mavirus in the presence of CroV. We propose that virophage integration and reactivation play an ecologically important role in regulating virus-mediated mortality of natural protist populations.

## Methods

### Host and virus strains

*C. roenbergensis* strain E4-10 was isolated from coastal waters near Yaquina Bay, OR, as described previously^16^. The cell suspension culture has since been continuously passaged approximately every 4 weeks in f/2 enriched natural or artificial seawater medium supplemented with 1-3 autoclaved wheat grains per 10 mL to stimulate bacterial growth. For infection experiments, cells were grown in f/2 enriched artificial seawater medium supplemented with 0.05% (w/v) Bacto^TM^ yeast extract (Becton, Dickinson and Company, Germany). For f/2 artificial seawater medium, the following sterile stock solutions were prepared: 75 g/L NaNO_3_; 5 g/L NaH_2_PO_4_; 1000x trace metal solution containing 4.36 g/L Na_2_EDTA x 2 H_2_O, 3.15 g/L FeCl_3_ x 6 H_2_O, 0.01 g/L CuSO_2_ x 5 H_2_O, 0.18 g/L MnCl_2_ x 4 H_2_O, 0.006 g/L Na_2_MoO_4_ x 2 H_2_O, 0.022 g/L ZnSO_4_ x 7 H_2_O, 0.01 g/L CoCl_2_ x 6 H_2_O; and a 50,000x vitamin solution containing 5 g/L thiamine-HCl, 25 mg/L biotin, and 25 mg/L cyanocobalamine. The vitamin solution was stored at −20°C, the other solutions at room temperature. To prepare 1 L of f/2 artificial seawater medium, 33 g of Red Sea Salt (Red Sea Meersalz, www.aquaristikshop.com) were dissolved in ultra-pure water (ELGA, Veolia Water Technologies, Germany), then 1 mL each of the 75 g/L NaNO_3_, 5 g/L NaH_2_PO_4_, and 1000x trace metal solutions as well as 20 µL of the 50,000x vitamin solution were added. After autoclaving, the medium was 0.22 µm filtered and stored at 4°C. Cultures were grown in flat-bottom 125 mL or 250 mL polycarbonate Erlenmeyer flasks (VWR, Germany) at 23°C in the dark.

The viruses used for infection experiments were Cafeteria roenbergensis virus (CroV) strain BVPW1 ^3,17^ and mavirus strain Spezl^2^.

### Viral infectivity assays

The infectivity of CroV was measured by end-point dilution assays and the statistical method by Reed and Muench^18^ was used to determine the 50% end point. The resulting cell culture infectious dose at which 50% of the cultures lysed (CCID_50_) was in good agreement with counts of SYBR-stained CroV particles by epifluorescent microscopy and also with gene copy numbers derived by quantitative PCR (qPCR). End-point dilution assays were carried out in 96-well plates with 200 µl of 1E+06 cells/mL exponentially growing host cells in f/2 medium + 0.05% (w/v) yeast extract per well. Each row (12 wells) was inoculated with a different dilution of CroV suspension (10 µL/well). Dilutions ranged from 1E-02 to 1E-09. The plates were stored at 23°C and analyzed after 6 days for cell lysis by microscopy. For mavirus, end-point dilution assays could not be employed because, in contrast to CroV, a productive mavirus infection does not result in cell lysis or cytopathic effects.

### Infection experiments

Typically, host cell suspension cultures were diluted daily to a cell density of (1-5)E+05 cells/mL with f/2 medium containing 0.05% (w/v) yeast extract, until the desired culture volumes were reached. On the day of infection, when the cells had reached a density of >1.0E+06 cells/mL, the cultures were diluted with f/2 medium containing 0.05% (w/v) yeast extract to a cell density of (5-7)E+05 cells/mL. Depending on the experiment, aliquots of 20 mL or 50 mL were dispensed in 125 mL or 250 mL polycarbonate flat-base Erlenmeyer flasks (Corning, Germany; through VWR International) and inoculated with virus-containing lysate or virus-free f/2 medium (for mock infections). The CroV inoculum varied between different infection experiments (see Supplemental Spreadsheet), according to the desired MOI and the titer of the CroV working stock, which was stored at 4°C and replaced every few months. Mock-infected cultures received an equal volume of f/2 medium. For testing culture supernatant from previous infection experiments for mavirus activity, 1 mL of the appropriate 0.1 µm pore-size filtered lysate were added to the flask immediately prior to the CroV inoculum. Cultures were incubated at 23°C in the dark. Cell concentrations were measured by staining a 10 µL aliquot of the suspension culture with 1 µL of Lugol’s Acid Iodine solution and counting the cells on a hemocytometer (Neubauer Improved Counting Chamber, VWR Germany). This method does not distinguish between live and dead cells and will also include cells that are already dead but have not lysed yet. Aliquots (200 µl) for DNA extraction were sampled at appropriate time points and were immediately frozen and stored at −20°C until further processing. All infections were carried out in triplicates, except those shown in Extended Data Fig. 1, which were done in single replicates.

### Isolation of *C. roenbergensis* strains E4-10P and E4-10M1

*C. roenbergensis* strain E4-10 was made clonal by repeated single-cell dilutions. Each well of a 96-well plate was filled with 200 µl of f/2 medium containing 0.01% (w/v) yeast extract. Then 1 µl of an E4-10 culture diluted to 300 cells/mL were added to each well, so that on average every 3^rd^ well received one cell. After 6 days at 23°C, wells were inspected for cell growth and positive samples were transferred to 20 mL of f/2 medium containing 0.05% (w/v) yeast extract. This procedure was repeated serially two more times. DNA from the final isolate was extracted and tested by qPCR to confirm the absence of mavirus. 20 mL cultures of the resulting E4-10P (parental) strain at 5E+05 cells/mL in f/2 medium containing 0.05% (w/v) yeast extract were then either mock-infected, infected with CroV at MOI=0.01, or co-infected with CroV (MOI=0.01) and mavirus (MOI≈1). Eight days post infection, the surviving cells from the co-infection were pelleted by centrifugation (5 min at 7,000 x g, 23°C), the pellets were resuspended in 50 mL f/2 medium and the centrifugation/dilution procedure was repeated 9 more times. The washed cells were then subjected to three consecutive rounds of single-cell dilution as described above. DNA was extracted from the resulting 66 clonal strains and tested by qPCR with mavirus-specific primers. The strain with the highest qPCR signal was named E4-10M1 (first mavirus-positive strain).

### Filtration Assay

Host strains E4-10P and E4-10M1 were either infected with CroV or mock-infected with f/2 medium. At 5 days p.i., when the CroV-infected cells had lysed, aliquots from the four different samples were passed through syringe filters of different nominal pore sizes, ranging from 5.0 µm to 0.1 µm, and DNA was extracted from 200 µl of each filtrate as well as from 200 µl of the unfiltered samples. The following syringe filters were used: 0.1 µm pore-size PVDF Millex (Millipore Merck, Ireland), 0.22 µm pore-size PES (TPP, Switzerland), 0.45 µm pore-size PES (TPP, Switzerland), 5.0 µm pore-size CN-S Whatman (Fisher Scientific GmbH, Germany). E4-10M1 cells were mechanically lysed by sonication with a Branson Sonifier 250 equipped with a microtip, duty cycle 50%, output setting 2. Two milliliter aliquots of an E4-10M1 suspension culture containing 1.4E+06 cells/mL were sonicated for 2x 30 sec with 30 sec incubation on ice in between. As a positive control, an E4-10P suspension culture was mixed with 0.1 µm-filtered reactivated mavirus to yield a final flagellate concentration of 1.8E+06 cells/mL. The sonicated and positive control samples were then filtered and processed as described above.

### DNA extraction and quantitative PCR

We used qPCR with the SYBR-related EvaGreen^TM^ dye to quantify viral DNA target sequences. Genomic DNA (gDNA) was extracted from 200 µL of suspension culture with the DNeasy 96 Blood & Tissue Kit (Qiagen, Hilden, Germany) following the manufacturer’s instructions for DNA purification of total DNA from cultured cells, with a single elution step in 100 µL of double-distilled (dd) H_2_O and storage at −20°C. DNA concentrations in the eluted samples typically ranged from 1 to 10 ng/µL, as measured on a NanoDrop 2000c spectrophotometer (Thermo Scientific, Germany). One microliter of gDNA was used as template in a 20 µL qPCR reaction containing 10 µL of 2X Fast-Plus EvaGreen^®^ Master Mix with low ROX dye (Biotium, Inc. via VWR, Germany), 10 pmol of each forward and reverse primer (see Table S3), and 8.8 µL of ddH_2_O. No-template controls (NTC) contained ddH_2_O instead of gDNA. Each qPCR reaction (sample, NTC, or standard) was carried out in technical duplicates, with individual replicates differing in their quantification cycles (*C*q) by about 0.5% on average (0.49% ± 0.43%, n=200). The limit of detection for this assay was ≈10 copies, which equates to ≈5000 copies per mL of suspension culture. The *C*q values of the NTC controls were consistently below the limit of detection. Thermal cycling was carried out in a Stratagene Mx3005P qPCR system (Agilent Technologies, Germany) with the following settings: 95°C for 5 min, 40 cycles of 95°C for 10 s followed by 60°C for 25 s and 72°C for 25 s, a single cycle of 72°C for 5 min, and a final dissociation curve was recorded from 50°C to 95°C. qPCR results were analyzed using MxPro™ qPCR software v4.10 (Stratagene, La Jolla, USA). The threshold fluorescence was set using the amplication-based option of MxPro™ software. Standard curves were calculated from a 10-fold dilution series that ranged from 10^1^ to 10^8^ molecules of a linearized pEX-A plasmid (Eurofins Genomics, Germany) carrying the fragment of the *MV18* MCP gene (GenBank Accession No: ADZ16417) that was amplified by primers Spezlq-PCR-5 and Spezl-qPCR-6 (Extended Data Table 3) for mavirus quantification, or gDNA extracted from a known amount of CroV particles, the concentration of which had been determined by epifluorescence microscopy. To directly compare the two different kinds of template DNA used for virus quantification, the linearized plasmid also contained the target sequence for the *crov283* gene (GenBank Accession No: ADO67316.1) that is amplified by primers CroV-qPCR-9 and CroV-qPCR-10 and used as an approximation for CroV genome copies. The resulting standard curves and *C*q values of the plasmid and gDNA templates were nearly identical, which implies that the quantification of mavirus using a plasmid-encoded target sequence is a valid approach. For mavirus quantification with primers Spezl-qPCR-5 and Spezl-qPCR-6, the R^2^ value for the standard curve was 0.996, the amplification efficiency was 109.7%, and the standard curve equation was Y=−3.109*log(x)+33.89. For CroV quantification with primers CroV-qPCR-9 and CroV-qPCR-10, the R^2^ value for the standard curve was 1.000, the amplification efficiency was 103.0%, and the standard curve equation was Y=−3.253*log(x)+34.77.

### PCR verification of an example mavirus integration site

The mavirus integration site shown in Fig. 1b was verified by PCR analysis and Sanger sequencing of the PCR products. Due to the difficulty of obtaining PCR products that were part host sequence with 70% GC content and part mavirus sequence with 30% GC content, several primers had to be tested under various PCR cycling conditions before the predicted products could be obtained. Primer sequences are listed in Extended Data Table 3. PCR amplifications were performed using 2 ng of genomic DNA template from strain E4-10P or E4-10M1 in a 25 µl reaction mix containing 5 µl Q5^®^ Reaction Buffer (NEB, Germany), 0.5 U of Q5^®^ High-Fidelity DNA Polymerase (NEB, Germany), 0.2 mM dNTPs and 0.5 µM of each primer. In addition, the PCR mixes to amplify the empty integration site with primers CrE_cont6-3 and CrE_cont6-6 (amplifying only host sequence with 70% GC content) contained 5 µl of Q5 High GC Enhancer solution. The PCRs were carried out in a TGradient thermocycler (Biometra, Germany) with the following cycling conditions: 30 s denaturation at 98°C; 35 cycles of 10 s denaturation at 98°C, 30 s annealing at 68°C (for primer pair CrE_cont6-3 & MaV37) or 69°C (for primer pairs MaV39 & CrE_cont6-6 and CrE_cont6-3 & CrE_cont6-6) and 1 min extension at 72°C; and a final 2 min extension at 72°C. For product analysis, 5 µl of each reaction were mixed with loading dye and pipetted on a 1% (w/v) agarose gel supplemented with GelRed. The marker lanes contained 0.5 µg of GeneRuler™ 1 kb DNA Ladder (Fermentas, Thermo-Fisher Scientific, USA). The gel was electrophoresed for 2 h at 70 V and visualized on a ChemiDoc™ MP Imaging System (BioRad, Germany). Cycling conditions for the PCR shown in Extended Data Figure 5 were: 45 s denaturation at 98°C; 35 cycles of 10 s denaturation at 98°C, 30 s annealing at 58°C (primer pairs MaV21F & MaV21R) and 1 min extension at 72°C; and a final 2 min extension at 72°C.

### RNA extraction and quantitative reverse-transcriptase PCR

Triplicate 50 mL cultures of strains E4-10P and E4-10M1 at an initial cell density of 6E+05 cells/mL were either mock-infected with f/2 medium or infected with CroV at an approximate MOI of 0.2. Aphidicolin-treated cultures were supplemented with 125 µl of a 2 mg/mL aphidicolin solution in DMSO (Sigma-Aldrich, Germany) for a final concentration of 5 µg/mL. Cycloheximide-treated cultures were supplemented with 37.5 µl of a 66.6 mg/mL cycloheximide solution in DMSO (Sigma-Aldrich, Germany) for a final concentration of 50 µg/mL. Cultures were incubated at 23°C. For extraction of total RNA, 1 mL aliquots were taken from each culture at 0 h p.i. and 24 h p.i. and centrifuged for 5 min at 10,000 x g, 21°C. The supernatants were discarded and the cell pellets were immediately flash-frozen in N_2_(l) and stored at −80°C until further use. RNA extraction was performed with the Qiagen RNeasy^®^ Mini Kit following the protocol for purification of total RNA from animal cells using spin technology. Cells were disrupted with QIAshredder homogenizer spin columns and an on-column DNase I digest was performed with the Qiagen RNase-Free DNase Set. RNA was eluted in 30 µl of 60°C warm RNase-free molecular biology grade water. The RNA was then treated with 1 µl TURBO DNase (2 U/µl) for 1 h at 37°C according to the manufacturer’s instructions (Ambion via ThermoFisher Scientific, Germany). RNA samples were analyzed for quantity and integrity on a Fragment Analyzer^TM^ capillary gel electrophoresis system (Advanced Analytical, USA) with the DNF-471 Standard Sensitivity RNA Analysis Kit. Six microliters of each RNA sample were then reverse transcribed into cDNA using the Qiagen QuantiTect^®^ Reverse Transcription Kit according to the manufacturer’s instructions. This protocol included an additional DNase treatment step and the reverse transcription reaction using a mix of random hexamers and oligo(dT) primers. Control reactions to test for gDNA contamination were done for all samples by omitting reverse transcriptase from the reaction mix. The cDNA was diluted twofold with RNase-free H_2_O and analyzed by qPCR with gene-specific primers. The qPCR reagents and conditions were the same as described above for genomic DNA qPCR. For data presentation purposes, any qPCR reactions that yielded no *C*q value after 40 PCR cycles were treated as *C*q=40. The no-template controls had an average *C*q value of 39.16 with a standard deviation of 2.20.

### Concentration, purification, and electron microscopy of reactivated mavirus particles

Five hundred milliliter cultures of strains E4-10P and E4-10M1 at 5E+05 cells/mL in 3 L polycarbonate Fernbach flasks were either mock-infected with f/2 medium or infected with CroV at an MOI of 0.02. Six replicates were prepared for a total volume of 3 L per condition (E4-10P or E4-10M1, mock-infected or CroV-infected). At 3 d p.i., the cultures were centrifuged for 40 min at 7000 x g and 4°C (F9 rotor, Sorvall Lynx centrifuge) and the supernatants were filtered on ice through a 0.2 µm PES Vivaflow 200 tangential flow filtration (TFF) unit (Sartorius via VWR, Germany). The filtrates were then concentrated on ice with a 100,000 MWCO PES Vivaflow 200 TFF unit to a final volume of ≈15 mL. The concentrates were passed through a 0.1 µm pore-size PVDF Millex syringe filter (Millipore Merck, Ireland) and analyzed on 1.1-1.5 g/mL continuous CsCl gradients. The CsCl gradients were prepared by underlayering 6.5 mL of 1.1 g/mL CsCl solution in 10 mM Tris-HCl, pH 8.0, 2 mM MgCl_2_ with an equal volume of 1.5 g/mL CsCl solution in 10 mM Tris-HCl, pH 8.0, 2 mM MgCl_2_ in a SW40 Ultra-Clear^TM^ centrifuge tube (Beckman Coulter, Germany). Tubes were capped and continuous gradients were generated on a Gradient Master (BioComp Instruments, Canada) with the following settings: tilt angle 81.5°, speed 35 rpm, duration 75 sec. After replacing 3.9 mL of solution from the top of the gradients with 4 mL of concentrated culture supernatants, the gradients were centrifuged for 24 h, 205,000 x g, 18°C using a SW40 rotor (Beckman Coulter, Germany) in a Beckman Optima™ ultracentrifuge. Bands in the gradients were visualized by illumination with an LED light source from the top of the gradient. One milliliter of gradient material from the mavirus band material (or equivalent positions of gradients were no such band was visible) were extracted with a syringe by puncturing the centrifuge tube with a 21G needle. The extracted band material was dialyzed for 24 h at 4°C in 3 mL dialysis cassettes (Pierce, 20 kDa cutoff) against 1 L of 10 mM Tris-HCl, pH 8.0, 2 mM MgCl_2_. After dialysis, each sample was diluted to 4 mL with 10 mM Tris-Cl, pH 8.0, 2 mM MgCl_2_ and centrifuged in Ultra-Clear^TM^ tubes (Beckman Coulter, Germany) in a SW60 rotor for 1 h, 100,000 x g, 18°C. The supernatant was discarded and the pellets were softened overnight at 4°C in 50 µl of 10 mM Tris-Cl, pH 8.0, 2 mM MgCl_2_ and then resuspended by pipetting. Aliquots (≈3 µL) of the concentrated samples were incubated for 2 min on Formvar/Carbon coated 75 mesh Cu grids (Plano GmbH, Germany) that had been hydrophilized by glow discharge. Grids were rinsed with ddH_2_O, stained for 90 sec with 1% uranyl acetate, and imaged on a Tecnai T20 electron microscope (FEI, USA) with an acceleration voltage of 200 kV.

### UV treatment of reactivated mavirus particles

A Stratalinker^®^ UV crosslinker 2400 (Stratagene) was used for irradiation of virus samples with UVC (λ=254 nm) light. Five hundred microliter drops of 0.1 µm-filtered reactivated mavirus suspension were pipetted on Parafilm and irradiated with a single dose of 500 J/m^2^ of UV-C light. The dose was monitored with a VLX 3W radiometer (Vilber-Lourmat). The irradiated virus suspension was then kept in the dark to prevent eventual light-induced DNA repair. Infection experiments were carried out as described above and cultures were incubated in the dark for the entire duration of the experiment. Samples for DNA extraction and qPCR analysis were taken and processed as described above.

### MiSeq and PacBio genome sequencing

Genomic DNA from 1E+09 cells each of the clonal *C. roenbergensis* strains E4-10P and E4-10M1 was isolated using the Qiagen Blood & Cell Culture DNA Midi Kit. The genomes were sequenced on an Illumina MiSeq platform (Illumina Inc., San Diego, USA) using the MiSeq reagent kit v3 at 2 x 300 bp read length configuration. The E4-10P genome was sequenced by GATC Biotech AG (Constance, Germany) with the standard MiSeq protocol. The E4-10M1 genome was prepared and sequenced at the Max Planck Genome Centre (Cologne, Germany) with NEBNext^®^ High-Fidelity 2X PCR Master Mix chemistry and a reduced number of enrichment PCR cycles (six) in order to reduce AT-bias. The total output was 6.8 Gbp and 4.5 Gbp for E4-10P and E4-10M1, respectively. Overall sequencing quality was assessed with FastQC v0.11.3. Reads were trimmed for low quality bases and adapter contamination using Trimmomatic v0.32^19^ and customized parameters (minimum phred score 20 in a 10 bp window, minimum length 75 bp, Illumina TruSeq3 reference adapter) resulting in 5.0 Gbp and 2.9 Gbp high quality paired-end sequences, respectively. We also sequenced genomic DNA of strains E4-10P and E4-10M1 on a Pacific Biosciences RS II platform (two SMRT cells each, Max Planck Genome Centre Cologne, Germany), which resulted in 0.52 Gbp and 1.30 Gbp of raw reads, respectively. The reads were extracted from the raw data files with DEXTRACTOR rev-844cc20 and general quality was assessed with FastQC v0.11.3.

### Read correction and assembly

Proovread v2.12^20^ was used for hybrid correction of the PacBio reads with the respective trimmed MiSeq read sets. Correction generated 423 Mbp (N50: 5994 bp) and 741 Mbp (N50: 7328 bp) of high accuracy long reads for E4-10P and E4-10M1, respectively. Reads were assembled into contigs with SPAdes v3.5.0^21^ using the dipspades.py module. Trimmed MiSeq reads were provided as paired-end libraries and corrected PacBio reads as single-end libraries. To account for structurally diverging sister chromosomes caused by asexual reproduction, the -expect-rearrangements flag was set. Assembly metrics were assessed with QUAST v2.3^22^. The E4-10P data set was assembled into 326 consensus contigs of at least 1000 bp, with a total assembly length of 40.3 Mbp and an N50 of 290 kbp. The E4-10M1 genome was assembled into 463 consensus contigs longer than 1000 bp, with a total assembly length of 31.4 Mbp and an N50 of 177 kbp.

### Proovread.cfg

~~~
#-- SI: proovread.cfg -----------------------#
‘seq-filter’ => {
‘--trim-win’ => “10,1”,
‘--min-length’ => 500,
},
‘sr-sampling’ => {
DEF => 0, # no sampling - entire sr-file
},
~~~

### Reference-guided assembly of the integrated mavirus genome

The E4-10M1 genome assembly was scanned for mavirus integration sites with blastn [NCBI BLAST v2.2.29+^23^]. The search returned one partial hit with 7000 bp and a few small hits with less than 600 bp alignment length. Additionally, partial hits were visualized and analyzed in context of the assembly graph structure using Bandage v0.4.2^24^. A full-length assembly of the potentially integrated mavirus genome sequence from the E4-10M1 set was generated through a reference guided assembly approach: Corrected PacBio reads of the E4-10M1 strain were aligned to the mavirus reference genome with blastn and strict settings (-evalue 10e-10-perc_identity 96). Matching reads longer than 1000 bp were extracted and assembled with SPAdes v3.5.0^21^ with the --only-assembler flag set.

### Detection/analysis of integration sites

Mavirus integration sites in the host genome were detected indirectly by identification of reads covering the junctions between a location in the *C. roenbergensis* genome and the terminal region of mavirus. In preparation, paired E4-10M1 MiSeq reads were merged with FLASH v1.2.11^25^ into longer single-end fragments to maximize the chances for unambiguous hits in subsequent mappings. The merged fragments as well as the corrected E4-10M1 PacBio reads were aligned to the revised TIR region of the mavirus genome with bwa mem [BWA v0.7.10-r984-dirty^26^] and SAMtools v1.1^27^. Fragments with a minimum alignment length of 30 bp and a minimum overlap of 10 bp at the TIR 5’ end were identified and extracted with a custom script. Due to the total length of 615/616 bp for the TIR, no merged MiSeq fragment spanned the entire TIR, and hence, no information about the strand-orientation of the mavirus core genome could be inferred from the MiSeq data. A read subset containing orientation information was generated by aligning extracted TIR-matching PacBio reads to the full mavirus genome and extracting end overlapping reads with a minimum alignment length of 650 bp. These reads spanned the entire TIR and extended into one side of the core region by at least 34 bp, thus yielding information about the orientation of the integrated element. The extracted mavirus end-overlapping MiSeq and PacBio reads were mapped with bwa mem onto the E4-10M1 genome assembly. Mapping locations of the reads were considered potential integration sites and have been further analyzed manually in a JBrowse^28^ genome browser instance, previously set up for the *C. roenbergensis* genome assemblies.

### Reconstruction of a mavirus integration site

Direct assembly of an integrated mavirus genome into the host genome was prevented by the diploid state of the *C. roenbergensis* genome and by the repetitive nature of the multiple mavirus integrations, which could not be properly resolved in assembly graph structures. Therefore, we manually reconstructed a contig comprising a mavirus integration site from the previously obtained integration site coordinate information and read evidence available in the MiSeq and PacBio data sets. For the reconstruction, we chose the predicted integration site at nucleotide position 118,064 on contig 5 (length: 208,205 bp). To validate the reconstructed sequence, MiSeq and corrected PacBio reads were mapped back against the artificial contig with bwa mem. Genomic features were annotated by mapping previously obtained host genome annotations (maker v2.31.8)^29^ and mavirus gene annotations (PROKKA v1.11 with custom mavirus database^30^) onto the new contig. Annotations were mapped with a custom script based on UCSC annotation lift-over strategies (<u>LiftOver_Howto, Minimal_Steps_For_LiftOver</u>) utilizing Kenttools v302^31^. Visualization of the annotated contig was generated with bio2svg v0.6.0.

### Ploidy assessment based on k-mer coverage frequency distribution

19-mer counts of the raw *C. roenbergensis* E4-10P Illumina MiSeq read data set were calculated with jellyfish v2.2.4^32^ in canonical representation and plotted with custom R scripts. Peak positions in Extended Data Fig. 2 were identified manually.

## Acknowledgements

This research was supported by the Max Planck Society. We are grateful to C. Suttle for access to host and virus strains, and to the Roscoff team for maintaining and distributing protist strains. We thank K.-A. Seifert, K. Fenzl and K. Barenhoff for technical assistance, U. Mersdorf for electron microscopy expertise, C. Roome for IT support, L. Czaja and the Max Planck Genome Centre in Cologne for bioinformatic assistance, S. Higgins for helpful suggestions, K. Haslinger and J. Reinstein for critical reading of the manuscript, and I. Schlichting for mentoring and support.

**Author Contributions** M.G.F. conceived the study, designed and performed experiments, collected, analyzed and interpreted data, and wrote the manuscript. T.H. corrected and assembled sequence data, and analyzed, interpreted and visualized data. Both authors revised and approved the manuscript.

## Author Information

*C. roenbergensis* strains E4-10P and E4-10M1 have been deposited in the Roscoff Culture Collection (strain numbers RCC 4624 and RCC 4625, respectively). The GenBank accession number for the reconstructed mavirus integration site of *C. roenbergensis* strain E4-10M1 is KU052222. The authors declare no conflict of interest.

## Extended Data

**Extended Data Figure 1:**
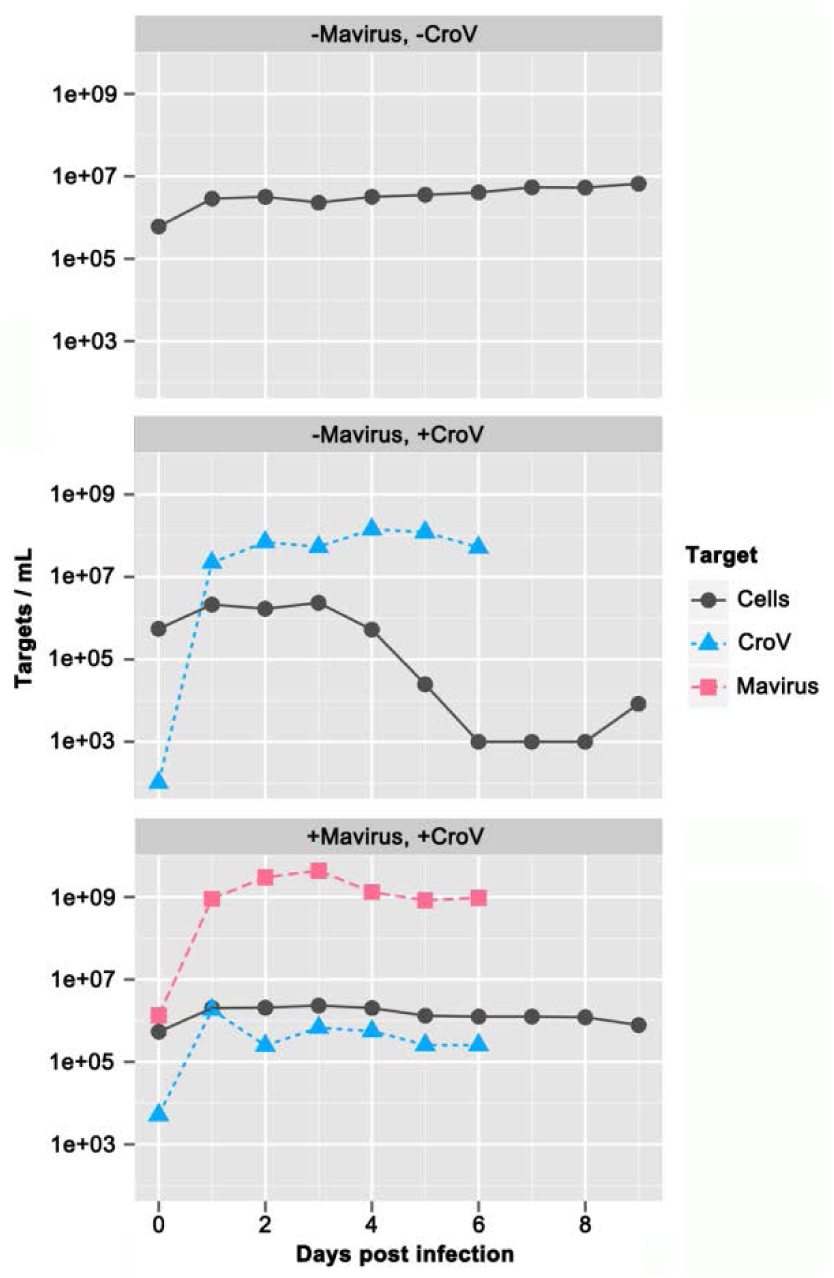
CroV-Mavirus co-infection experiment of *C. roenbergensis* strain E4-10P to generate the mavirus-carrying strain E4-10M1. *C. roenbergensis* strain E4-10P was either mock-infected, infected with CroV, or co-infected with CroV and mavirus. Cell densities are based on microscopy counts and were monitored for 8 days, viral numbers were monitored for 6 days and are derived from qPCR data assaying short amplicons of the mavirus *MV18* gene (MCP) and the *crov283* gene (D11-like transcription factor), respectively. The detection limit for both methods was ≈1E+03/mL. These experiments were carried out in single copies. The mavirus-positive host strain E4-10M1 was isolated from the pool of surviving cells in the +mavirus, +CroV infection.

**Extended Data Figure 2:**
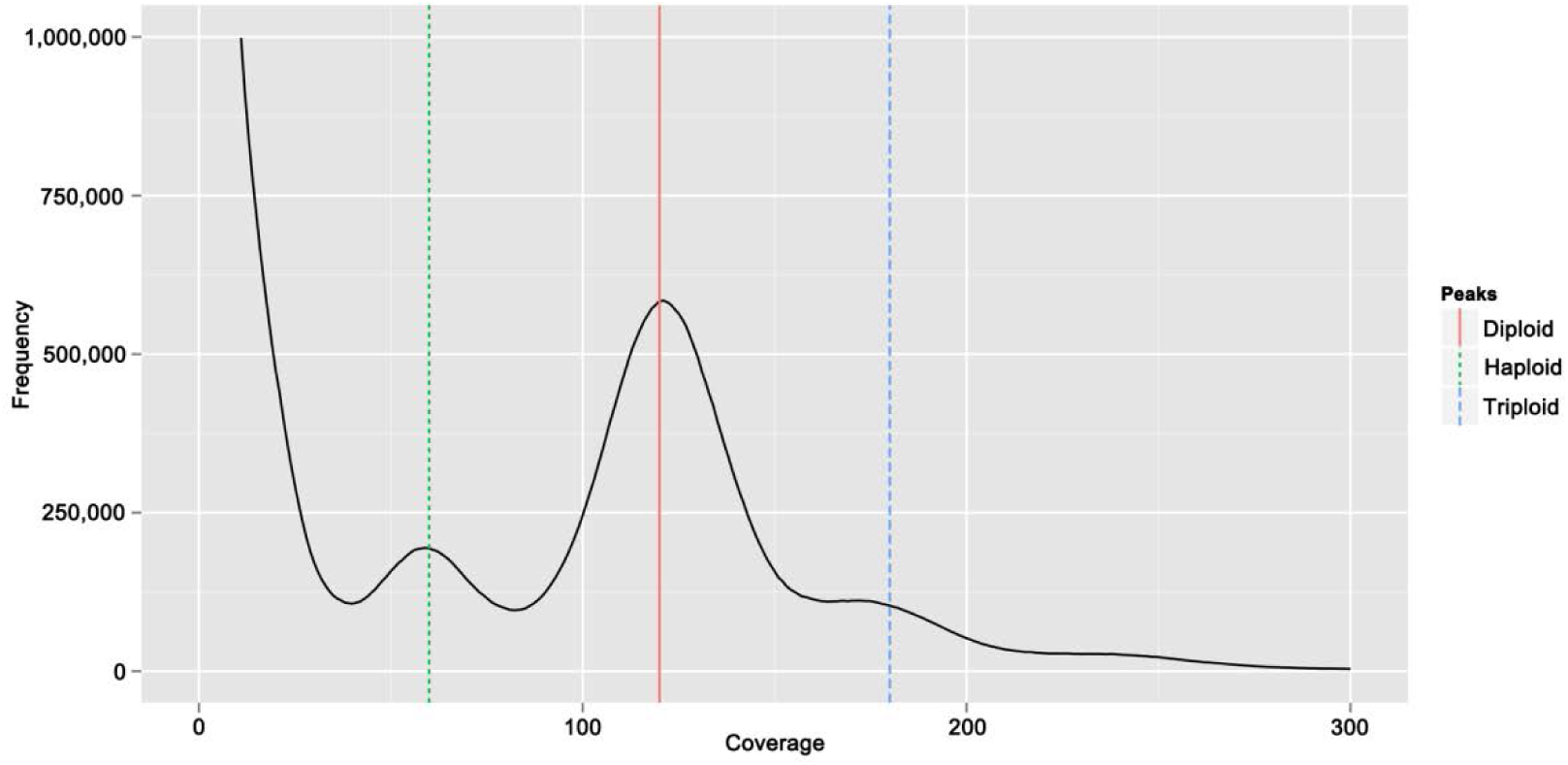
K-mer frequency distribution of the diploid genome of *Cafeteria roenbergensis* strain E4-10P. Frequency distribution of *in silico* generated random 19-mers in the genomic read set of E4-10P. The distribution exhibits a major peak at 120X coverage (solid red line) corresponding to the majority of homozygous k-mers of the underlying diploid genome, a smaller peak at half the diploid coverage (60X, dotted green line) comprising haplotype-specific k-mers, and a weak third peak at three times the haploid coverage (180X, dashed blue line) indicating a primarly diploid, partially triploid genome structure. Low coverage k-mers derive from sequencing errors and bacterial contamination.

**Extended Data Figure 3:**
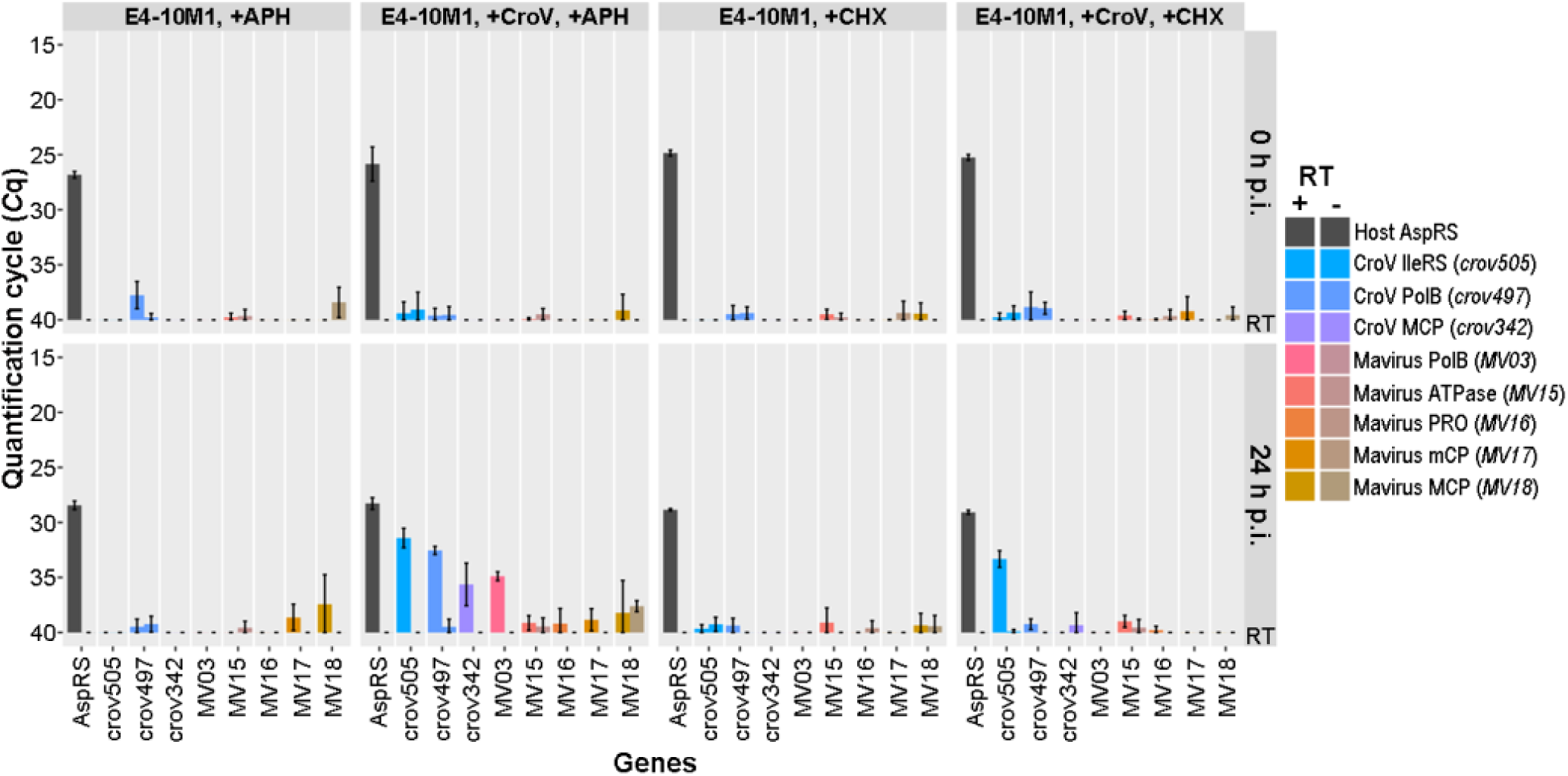
Gene expression of the endogenous mavirus genome is inhibited by cycloheximide and aphidicolin. Selected cellular and viral transcripts isolated at 0 and 24 h p.i. from mock-infected or CroV-infected E4-10P and E4-10M1 cultures in the presence of 5 µg/mL aphidicolin or 50 µg/mL cycloheximide with were quantified by qRT-PCR. Shown are the average quantification cycle (Cq) values of three independent experiments with error bars representing ± SD. The following genes were assayed: host AspRS, *C. roenbergensis* E4-10 aspartyl-tRNA synthetase; crov342, CroV major capsid protein; crov497, CroV DNA polymerase B; crov505, CroV isoleucyl-tRNA synthetase; MV03, mavirus DNA polymerase B; MV15, mavirus genome-packaging ATPase; MV16, mavirus maturation protease; MV17, mavirus minor capsid protein; MV18, mavirus major capsid protein. Cq values of the reverse transcriptase-negative (-RT) reactions are shown directly to the right of the respective +RT results. Accession numbers are listed in Extended Data Table 3. See also Supplemental Spreadsheet.

**Extended Data Figure 4:**
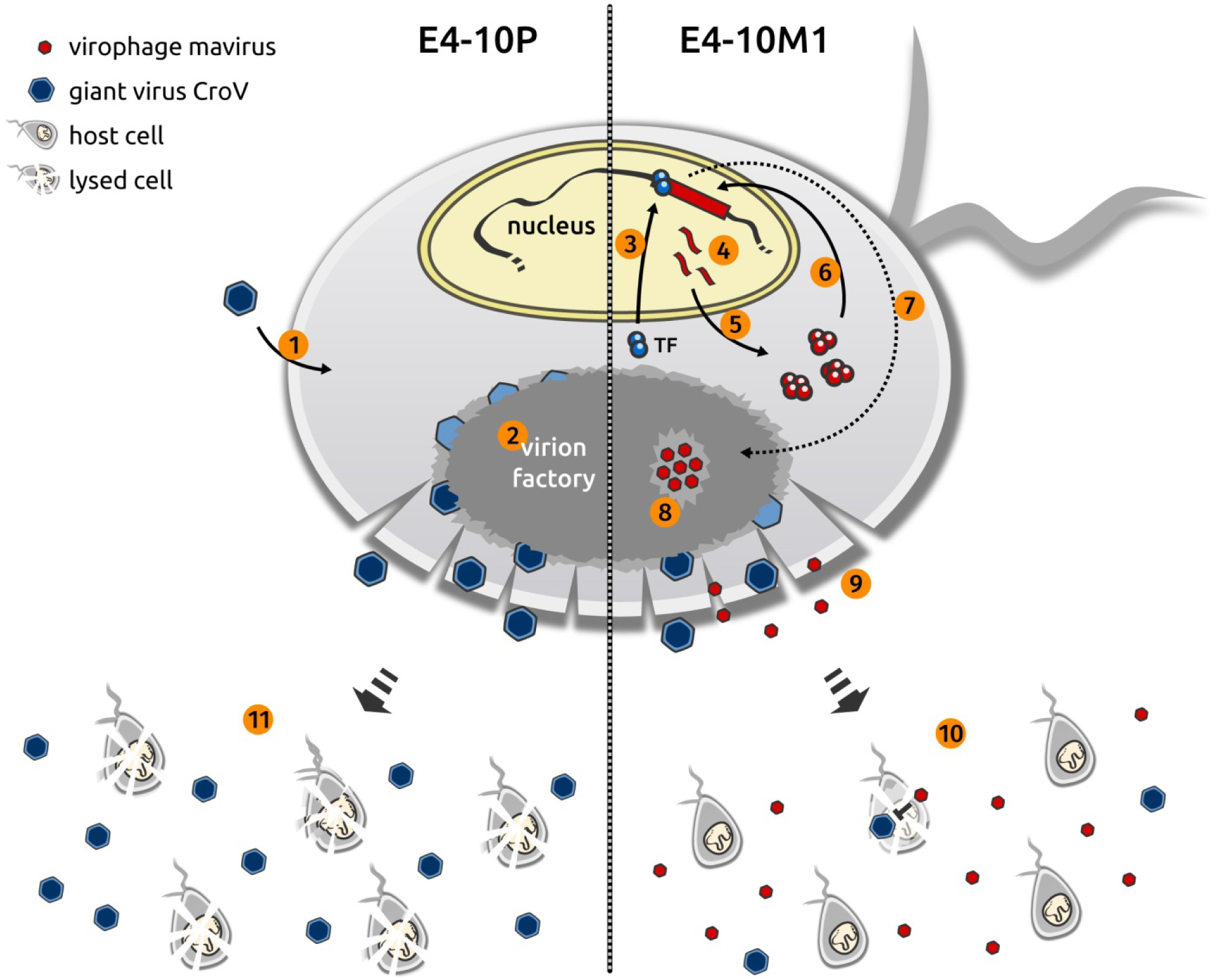
Hypothesis for CroV-induced reactivation of endogenous mavirus. Shown is a schematic *C. roenbergensis* cell displaying selected events of a CroV infection cycle in strains E4-10P (left) and E4-10M1 (right). Following CroV entry (1), the virion factory forms in the cytoplasm. At the onset of late phase, a CroV-encoded transcription factor (TF) recognizing the late CroV promoter motif is synthesized (2). In E4-10M1 cells, the late TF enters the nucleus (3), binds the mavirus promoter sequences and activates gene expression of the provirophage (4). Mavirus-specific transcripts are exported and translated (5) and some of the mavirus proteins return to the nucleus to excise or replicate the provirophage genome (6). The mavirus genome then translocates to the CroV factory (7), where genome replication, particle assembly, and genome packaging occur (8). Cell lysis releases the newly synthesized CroV and mavirus particles (9) and the reactivated virophages inhibit further spread of the CroV infection in co-infected cells, leading to enhanced survival of the host population (10). In contrast, CroV infection of an E4-10P cell does not induce a virophage response and CroV continues to infect other host populations (11).

**Extended Data Figure 5:**
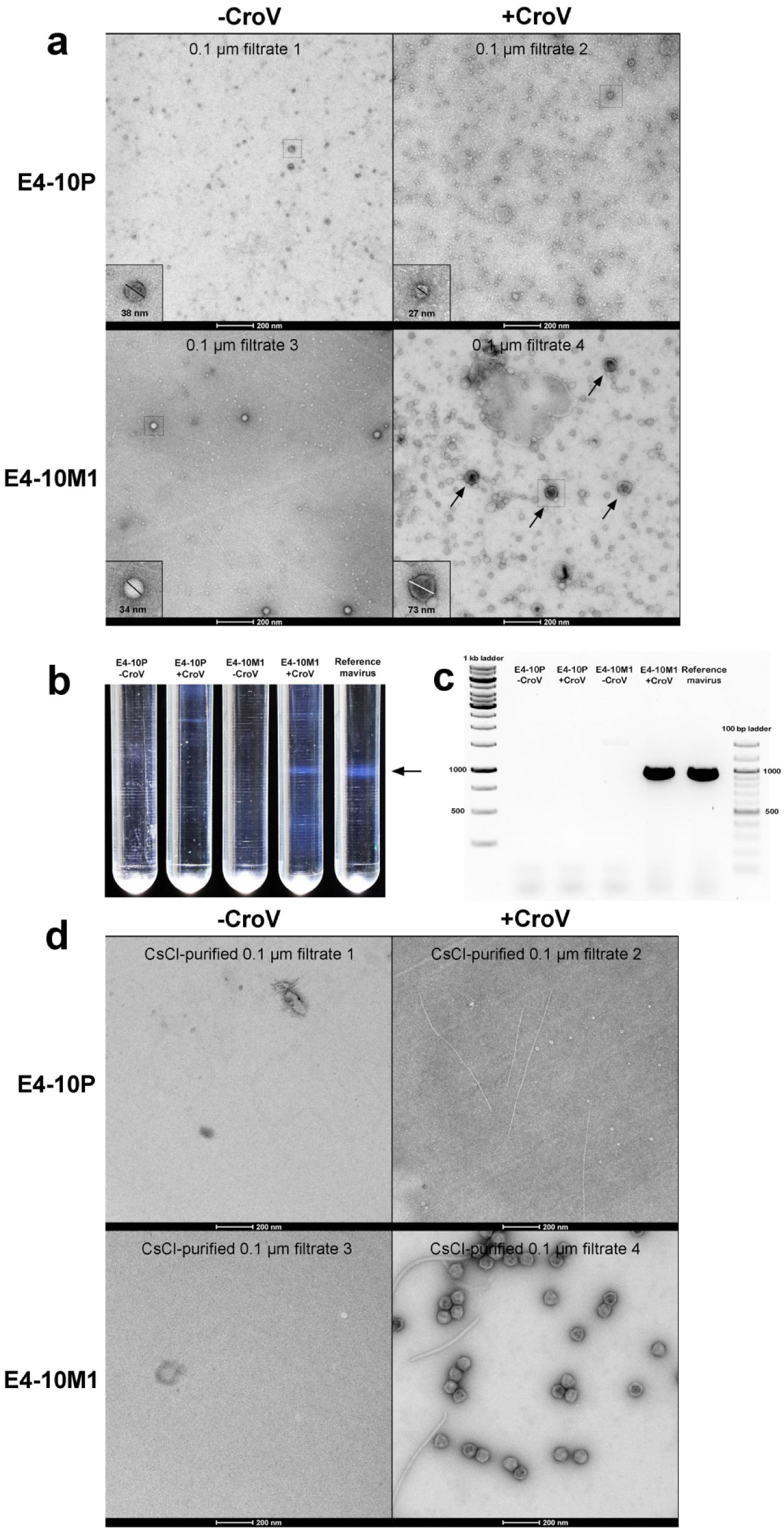
Purification and characterization of reactivated mavirus particles. **a**, Three-liter cultures of CroV- or mock-infected E4-10P and E4-10M1 cultures were concentrated 200-fold and stained with uranyl acetate for electron microscopy. Representative particles from each filtrate are boxed and shown at higher magnification. Mavirus-like particles are marked by arrows. Filtrate numbers refer to the infections shown in Fig. 3a. **b**, The concentrated samples were analyzed on 1.1-1.5 g/mL linear CsCl density gradients. A concentrated sample of reference mavirus was run in parallel and yielded a band at ≈1.29 g/mL CsCl (arrow). Only the CroV-infected E4-10M1 culture produced a band at a similar density. **c**, PCR analysis of band material extracted from the CsCl gradients shown in **b**) at a density of ≈1.29 g/mL CsCl. Primers MaV21F & MaV21R were used to generate a 956 bp long product of the *MV19* gene. **d**, Material from the 1.29 g/mL CsCl band or from equivalent positions was extracted from the gradients and visualized by negative-stain electron microscopy. Only the CroV-infected E4-10M1 culture contained mavirus-like particles.

**Extended Data Figure 6:**
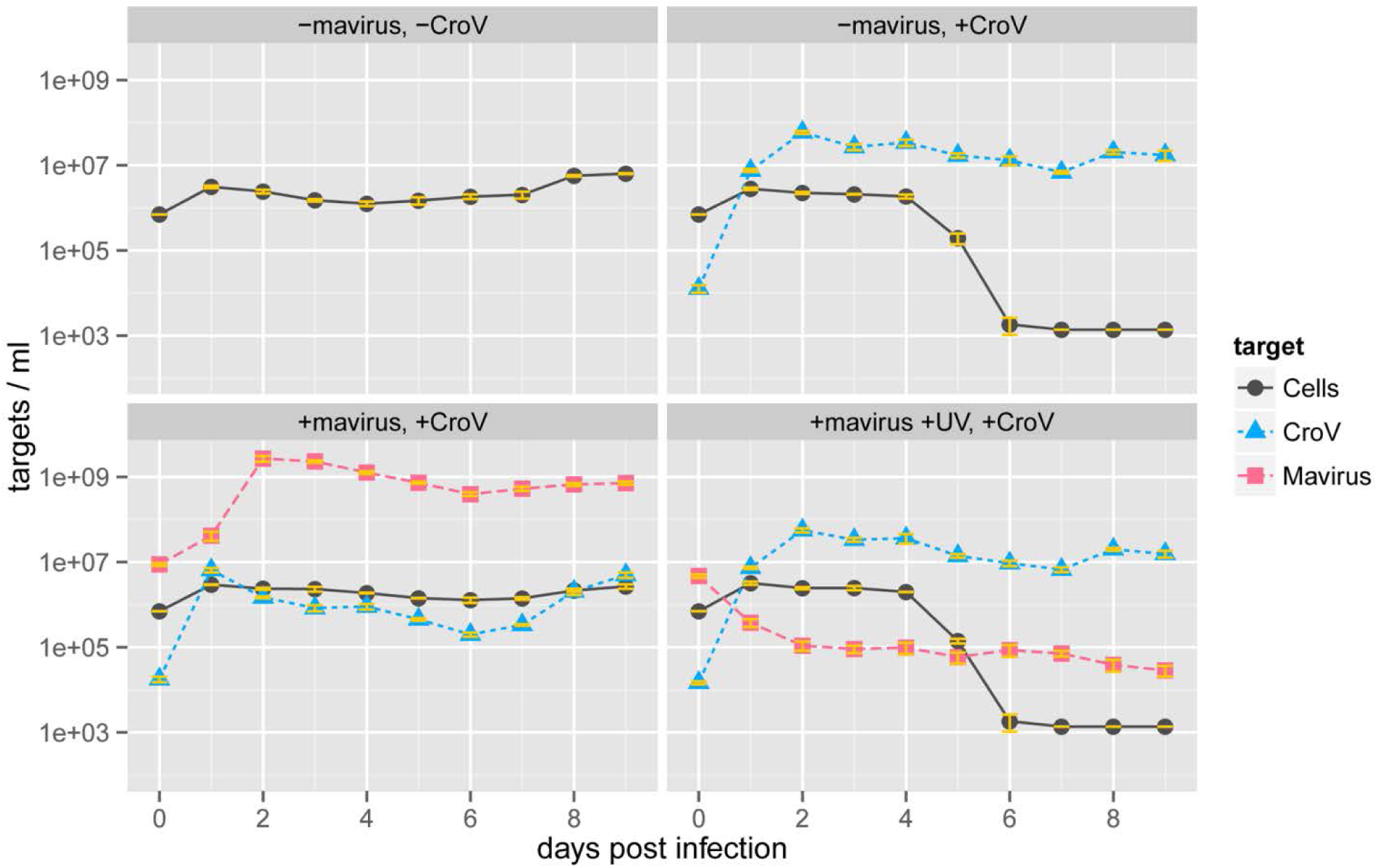
UV treatment abolishes infectivity of reactivated mavirus. Reactivated mavirus contained in filtrate 4 from the infection experiments shown in Fig. 3a was irradiated with 500 J/m^2^ of UV-C light (λ=254 nm). UV-treated and untreated mavirus suspensions were tested for infectivity by co-infection of host strain E4-10P with CroV. As shown in the lower right panel, reactivated mavirus treated with UV light was no longer able to replicate in the presence of CroV. The data are shown as the mean of biological triplicates ± SD. The numerical data of the individual replicates for these and all other infection experiments are listed in the Supplemental Spreadsheet.

**Extended Data Table 1:** Cell and virus concentrations in different size fractions of mock-infected and CroV-infected E4-10P and E4-10M1 populations. Uninfected host cultures or CroV-infected cultures after cell lysis were passed through syringe filters of various nominal pore sizes. As controls, E4-10P cells were spiked with mavirus particles immediately prior to filtration, and uninfected E4-10M1 cells were mechanically lysed by sonication and then filtered. DNA extracted from identical volumes of each filtrate was used as template in qPCR assays with mavirus- and CroV-specific primers. Cell concentrations were determined by microscopy counts. Shown are the average values of three independent experiments with error bars representing ± SD. *C. roenbergensis* cells are 5-10 µm in diameter, CroV particles are 300 nm in diameter, mavirus particles are 75 nm in diameter. BDL, below detection limit (≈1E+03 cells or viruses per mL); CN, cellulose nitrate; PES, polyethersulfone; PVDF, polyvinylidene difluoride.

| Condition | Host strain | Target | Unfiltered | 5.0 µm filtrate (CN) | 0.45 µm filtrate (PES) | 0.22 µm filtrate (PES) | 0.1 µm filtrate (PVDF) |
| --- | --- | --- | --- | --- | --- | --- | --- |
| Uninfected | E4-10P | Cells/mL | (1.53±0.18)E+06 | (9.20±4.20)E+03 | BDL | BDL | BDL |
|  |  | Mavirus/mL | BDL | BDL | BDL | BDL | BDL |
|  |  | CroV/mL | BDL | BDL | BDL | BDL | BDL |
|  | E4-10M1 | Cells/mL | (1.39±0.14)E+06 | (5.50±0.00)E+03 | BDL | BDL | BDL |
|  |  | Mavirus/mL | (4.03±0.61)E+06 | (1.04±0.46)E+04 | BDL | BDL | BDL |
|  |  | CroV/mL | BDL | BDL | BDL | BDL | BDL |
| CroV-infected | E4-10P | Cells/mL | BDL | BDL | BDL | BDL | BDL |
|  |  | Mavirus/mL | BDL | BDL | BDL | BDL | BDL |
|  |  | CroV/mL | (5.40±0.87)E+07 | (1.35±0.10)E+06 | (2.34±1.00)E+06 | (3.11±0.47)E+04 | (9.55±4.00)E+04 |
|  | E4-10M1 | Cells/mL | BDL | BDL | BDL | BDL | BDL |
|  |  | Mavirus/mL | (1.89±0.17)E+09 | (1.25±0.05)E+09 | (9.90±0.53)E+08 | (8.97±0.71)E+08 | (4.76±0.59)E+08 |
|  |  | CroV/mL | (1.41±0.12)E+08 | (3.03±0.15)E+07 | (1.00±0.03)E+07 | (1.48±0.48)E+04 | (1.18±0.20)E+04 |
| Mavirus-spiked | E4-10P | Cells/mL | (1.83±0.19)E+06 | (2.29±0.64)E+04 | BDL | BDL | BDL |
|  |  | Mavirus/mL | (1.22±0.02)E+09 | (7.37±4.67)E+08 | (7.64±4.52)E+08 | (9.74±0.72)E+08 | (7.23±0.14)E+08 |
|  |  | CroV/mL | BDL | BDL | BDL | BDL | BDL |
| Mechanically lysed | E4-10M1 | Cells/mL | (1.83±1.41)E+04 | BDL | BDL | BDL | BDL |
|  |  | Mavirus/mL | (1.24±0.05)E+07 | (1.15±0.04)E+07 | (1.16±0.02)E+07 | (1.17±0.05)E+07 | (1.17±0.10)E+07 |
|  |  | CroV/mL | BDL | BDL | BDL | BDL | BDL |

**Extended Data Table 2:**
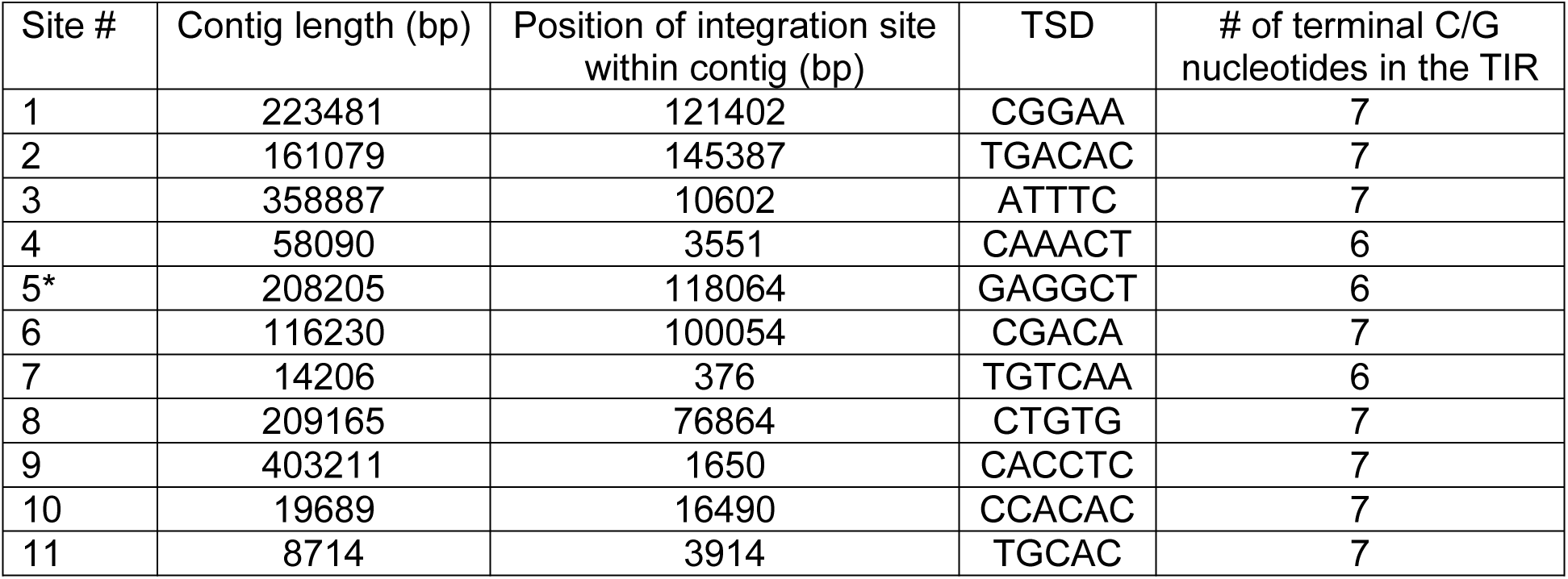
Details on the 11 bioinformatically well-supported mavirus integration sites in *C. roenbergensis* strain E4-10M1. The integration site described in detail in Fig. 1b is marked with an asterisk.

**Extended Data Table 3:**
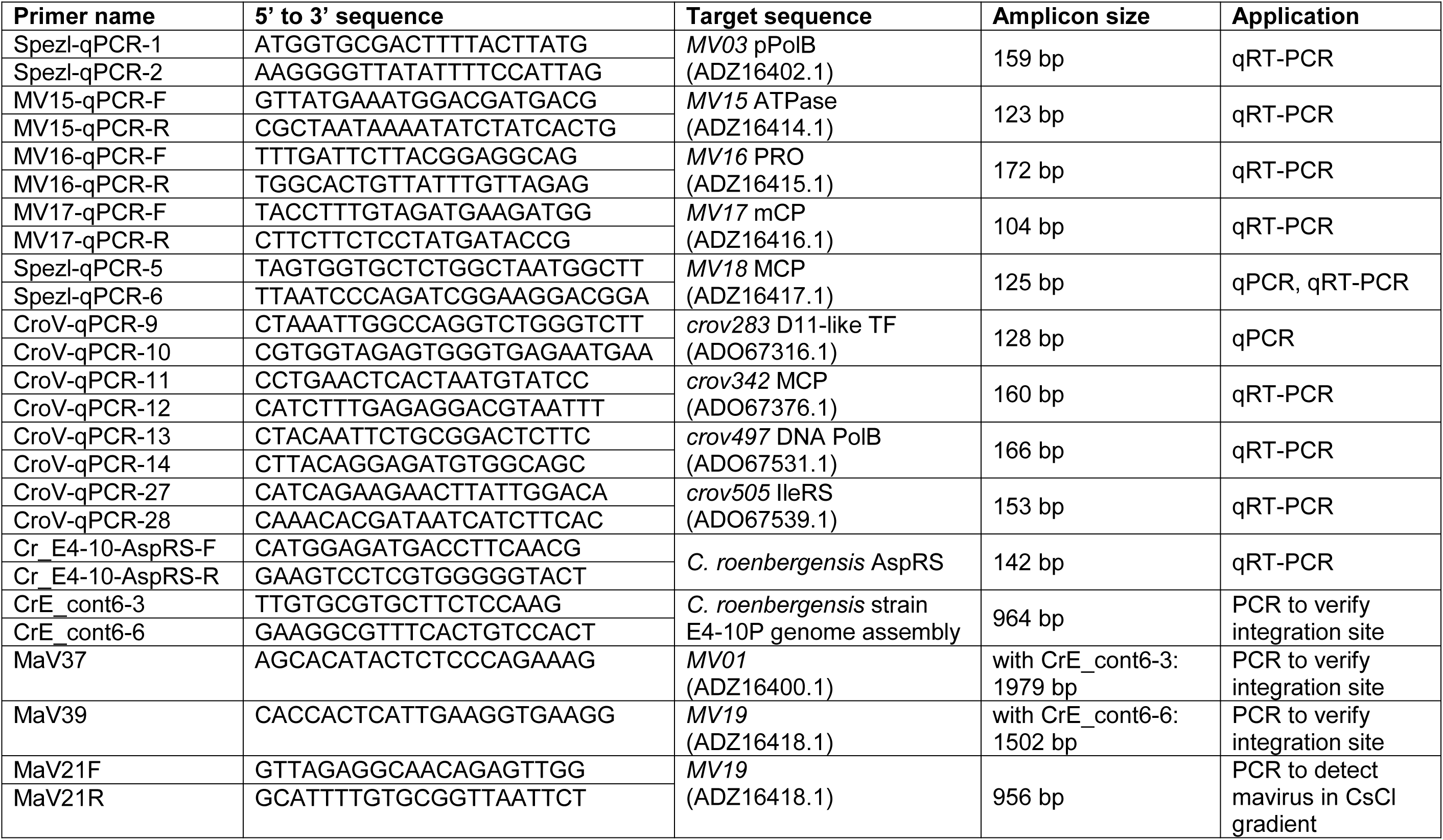
PCR oligonucleotide primers used in this study. GenBank accession numbers are listed where available.

